# Liposomes Under Real Cell Conditions Behave Like Real Cells with a Single Pulsed Electric Field

**DOI:** 10.1101/860841

**Authors:** Gen Urabe, Masaharu Shimada, Takumi Ogata, Sunao Katsuki

## Abstract

Liposomes are widely assumed to present a straightforward physical model of cells. However, almost all previous liposome experiments with pulsed electric fields (PEFs) have been conducted in low-conductivity liquids, a condition that differs significantly from that of cells in medium. Here, we prepared liposomes consisting of soy bean lecithin and cholesterol, at a molar ratio of 1:1, in higher-conductivity liquid that approximated the conditions of red blood cells in phosphate-buffered saline, with inner and outer liquid conductivities of 0.6 and 1.6 S/m, respectively. We found that a single 1.1 kV/cm, 400 μs PEF promoted cell-like spontaneous division of liposomes.

## Introduction

Bioelectrics, which studies the relationship between electricity and biology, has been the subject of both basic and applied research of note [1–9]. Electropores induced on plasma membranes may trigger responses in cells and tissues to pulsed electric fields (PEFs) [10–17]. Numerous papers have attempted to validate this hypothesis, applying pulses to liposomes composed of artificial lipid membranes [1,18–20]. However, few studies have managed to completely mimic the phenomenon in cells. In the case of cells, plasmid DNA reportedly accumulates on the cell membrane facing the cathode side, and subsequently, actin patches appear on the surface, after which plasmid DNA is taken into the cells [3,21]. In a study using liposomes, plasmid DNA traversed the lipid membrane without accumulating [22]. Rols et al. assumed actin as the source of the difference and suggested conducting a similar experiment using actin-containing liposomes.

Little dye was seen in liposomes without actin, but liposomes with actin proceeded taking dye more than 146 s, and the latter reaction was similar to those of cells under a PEF [23]. However, those liposomes were prepared in liquid with much lower conductivity than is seen in cytoplasm and extracellular liquid (phosphate-buffered saline [PBS]) [24–27]. Because liquid conductivity influences substance influx through electropores on cells [28–30] and low-conductivity conditions differ from those of cell experiments, experiments that approximate real cytoplasm and media will require adjustment of the conditions. Furthermore, although the liposomes exhibited accurate lipid molar fractions, it is unlikely that their composition was similar to that of real cell membranes. As membrane composition can influence reactions to PEFs, liposome compositions that are closer to those of a real plasma membrane are necessary. To that end, we adopted an emulsion method that made it possible to adjust the conductivities of the interior and exterior liquids of the liposomes. We applied PEFs to liposomes with inner- and outer-liquid conductivities comparable to those of red blood cells (0.6 S/m) and PBS (1.6 S/m) and whose membranes were composed of soybean lecithin and cholesterol. We adjusted the conductivity of liposomes’ inner liquid to that of red blood cells because their value had already been reported, and most previous simulation studies preferred this value [31]. Because soybean lecithin is not a purified lipid, but an extract, we assumed that the composition of soybean lecithin was close to that of real cell membranes. Although the actual components are uncertain, lecithin is composed mainly of phosphatidylcholine (PC), which is a main component of cell membranes. We also applied egg lecithin to the process but were unavailable to produce liposomes.

## Materials and methods

### Materials

For liposome preparation, we purchased soy bean lecithin, cholesterol, liquid paraffin, 1,2-dioleoyl-sn-glycero-3-phosphocholine (DOPC), 1- palmitoyl- 2- oleoyl- sn- glycero- 3-phosphocholine (POPC), chloroform, glucose, and sucrose from Wako and Texas Red- 1,2- dihexadecanoyl- sn- glycero- 3- phosphoethanolamine (DHPE) from Takara. For the liposome outer liquid, we used PBS(-) (from Wako), and for the liposome inner liquid, we mixed PBS(-) with 308 mM glucose solution at a 15:26 mL volume ratio. The theoretical osmotic pressure of PBS(-) was 306 mOsm/L. The osmotic pressure of the 308 mM glucose solution was theoretically 308 mOsm/L. Conductivities of the inner and outer liquids were 0.64 and 1.64 S/m, respectively. Cholesterol, DOPC, and POPC were dissolved in chloroform at 180 mM and Tex Red-DHPE at 0.9 mM. All solutions were stored at a temperature of −20°C.

### Liposome preparation

#### Lecithin liposome

Liposome production followed the water-in-oil emulsion method [32]. We estimated the soy bean lecithin molar weight at 758.06 and then dissolved 0.53 mg of soy lecithin in 500 μL of liquid paraffin at 80°C by vortex (1.4 mM). After storing the sample at a temperature of 80°C for 10 min with the cap of the microtube left open, we added 50 μL of the inner liquid and vortexed it for 10 s and then immediately put the sample on ice for 10 min to stabilize the emulsions. Next, 400 μL of the emulsion solution was put on 400 μL of the outer liquid, which had been on ice. The volume ratio of the emulsion solution and the outer liquid was 400:400. After treating the sample under centrifugation at 18,000 *g* and a temperature of 4 °C for 5 min, we disposed of as much of the supernatant as possible and extracted the sediments, which was the aggregation of liposomes.

### Lecithin-cholesterol liposomes and fluorescent liposomes

To produce cholesterol-containing lecithin liposomes (at a molar ratio of lecithin/cholesterol = 1:1), we added 4 μL of cholesterol solution to a heated liquid paraffin-lecithin solution at a temperature of 80°C and then mixed them by vortex to adjust the cholesterol concentration to 1.4 mM. For a lecithin/cholesterol molar ratio of 1:1.5 and 2:1 liposomes, the amounts of added cholesterol solution were 6 and 2 μL. For fluorescent labeling, 2 μL of Tex Red-DHPE was added simultaneously to make a final concentration of 3.6 μM (with a molar ratio of 0.13 mol%). The sample was stored at a temperature of 80°C for 10 min with the cap of the microtube left open, and the rest of the protocol was the same as that of lecithin liposome.

### DOPC liposomes and POPC liposomes

To produce DOPC liposomes, DOPC and cholesterol liquids were mixed in 500 μL of liquid paraffin at the following ratios: 4:0 for DOPC-only liposomes, 4:4 for DOPC/cholesterol = 1:1 (molar ratio), 2:4 for DOPC/cholesterol = 1:2 (molar ratio), and 2:6 for DOPC/cholesterol = 1:3 (molar ratio). In the case of POPC, the POPC and cholesterol liquids were mixed at the following ratios; 4:0 for POPC-only liposomes and 4:4 for POPC/cholesterol = 1:1 (molar ratio). The sample was stored at a temperature of 80°C for 10 min with the cap of the microtube left open, and the rest of the protocol was the same as that of lecithin liposome

### Calcium-ion influx detection

#### The same osmotic pressure between inner and outer liquid

First, 50 μg of Fluo-8(R) sodium salt (Cosmo Bio) was dissolved in 62.5 μL of Milli-Q, and 1 μL of the solution was diluted in 50 μL of the inner liquid for a final Fluo-8 concentration of 20 μM. For the calcium-ion flow experiment, liposomes were prepared with the Fluo-8-containing inner liquid. After preparing the liposome sample, a solution of D-PBS(+) preparation with a Ca and Mg reagent solution 100× (Nacalai Tesque) was added at 1 % (v/v) to mix the calcium ions in the outer liquid.

#### Different osmotic pressures between inner and outer liquid

To strengthen calcium influx, we set the outer-liquid osmotic pressure at half of that of the inner liquid. The inner-liquid composition was the same as that used to text Ca-ion influx, but in the case of the outer liquid, the PBS (-) was diluted twice with Milli-Q. The rest of the protocol was the same as that of no osmotic pressure difference.

### PEF application system and microscopy

We used two slices of a platinum plate, which are 0.2-mm thick and 2.7-mm wide, as electrodes. The electrodes were fixed with a 640-μm gap on a glass slide and connected to the PEF generator. We set the glass slide–electrode device on a fluorescent microscope (Leica, DMi8) combined with a digital camera (Canon, EOS 8000D). Movies were recorded at 67 frames per second. The details of microscopy are provided in a previous paper [33].

## Results

We referred to HeLa cells, which were familiar to us, for the lipid molar ratio of cell membranes, facilitating a comparison of liposome and cell results. As PC and cholesterol constitute primarily of HeLa cell membranes, their molar ratio was 1:1, and other lipids represented only minor parts [34,35], we mixed lecithin and cholesterol at a molar proportion of 1:1.

### Liposomes divided spontaneously

When we applied a single 1.1 kV/cm, 400 μs PEF to the liposomes, the liposomes divided spontaneously (Fig. 1A, B, and C). To determine the optimal PEF condition to induce division, we scanned the pulse duration and electric field intensity, showing that a 400 or 500 μs, 1.1 kV/cm PEF appeared to have the potential to induce division (Table. 1). A PEF with identical energy but a shorter pulse did not promote self-division (Table. 2). Because a single 1.1 kV/cm, 400 μs PEF can theoretically increase the PBS temperature by 2°C, the thermal influence may be small. These results suggest that energy was not an important factor in the division.

**Figure 1.**
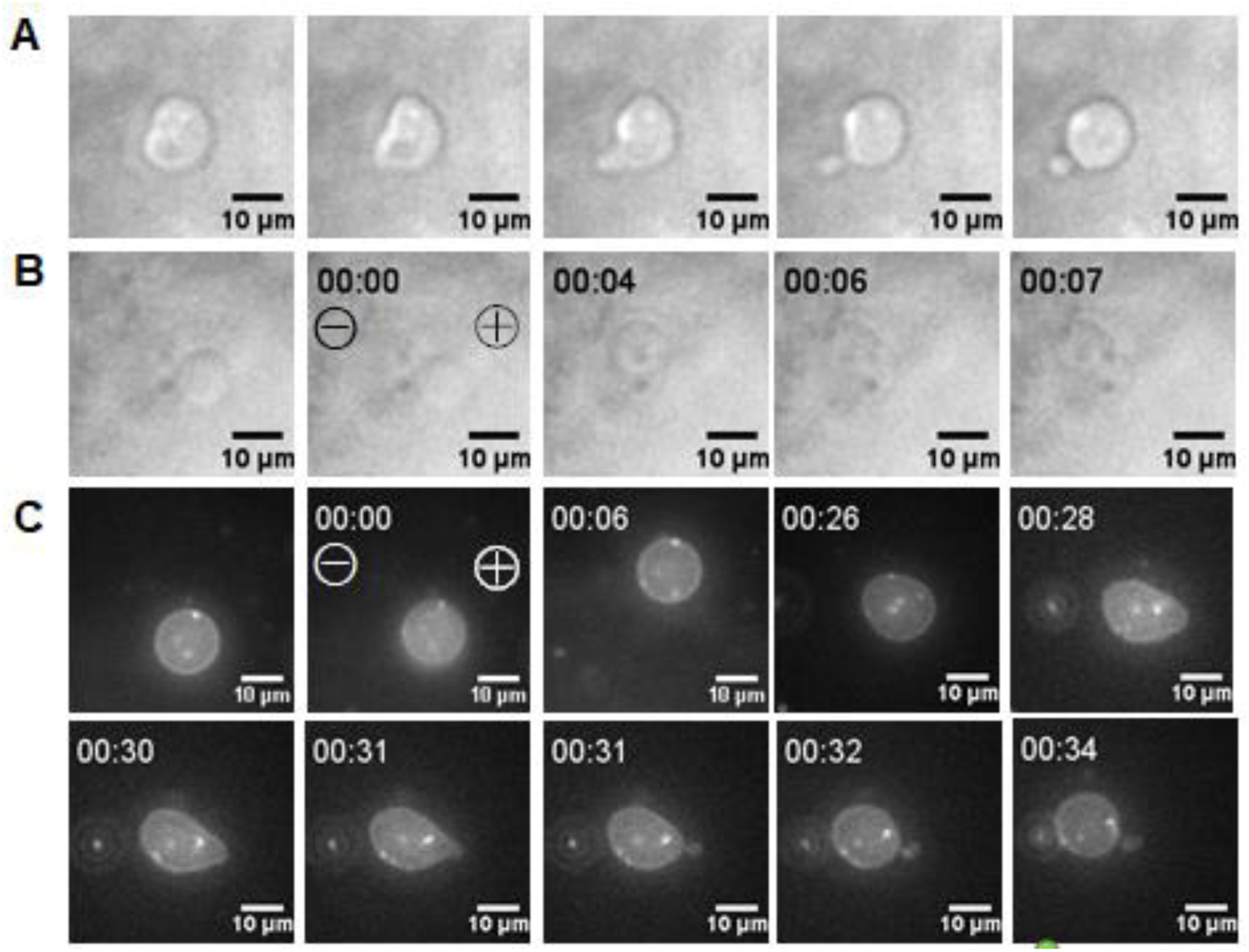
A) Spontaneous liposome divisions and PEF conditions. B) Lecithin/cholesterol = 1:1 (molar ratio) liposomes divided spontaneously after a 1.1 kV/cm, 400 μs single PEF application. As we did not measure the time, there is no time display in series A. All time displays are in seconds. “00:00” = timing of PEF application. C. Liposomes with lecithin/cholesterol/texas red-DHPE = 1.4:1.4:0.0036 mM. ⊕ = the anode side of the electrode; ⊖ = the cathode side of the electrode.

**Table 1.** Various PEF duration vs. whether the liposome division happened. Electric field was 1.1 kV/cm.

| Pulse duration / $\mu\text{s}$ | Division |
| --- | --- |
| 100 | × |
| 200 | × |
| 300 | △ |
| 400 | $\circ$ |
| 500 | $\circ$ |

$\circ$ = liposome division occurred almost every time; △ = division occurred some of the time; × = division did not occur.

**Table 2.** Two different PEF conditions with the same energy of 1.1 kV/cm PEF (400 μs). Both conditions did not trigger liposome division.

| Electric field (kV/cm) | Pulse duration ($\mu\text{s}$) | Division |
| --- | --- | --- |
| 6.9 | 10 | × |
| 21.9 | 1 | Discharged |

× indicates no liposome division

### Do cholesterol and lecithin have division ability?

As liposomes without cholesterol did not divide, it is possible that cholesterol has the potential to induce division. Varying the molar ratios of cholesterol, liposomes with more than 50% of cholesterol divided (Fig. 2A, B, C, and D).

**Figure 2.**
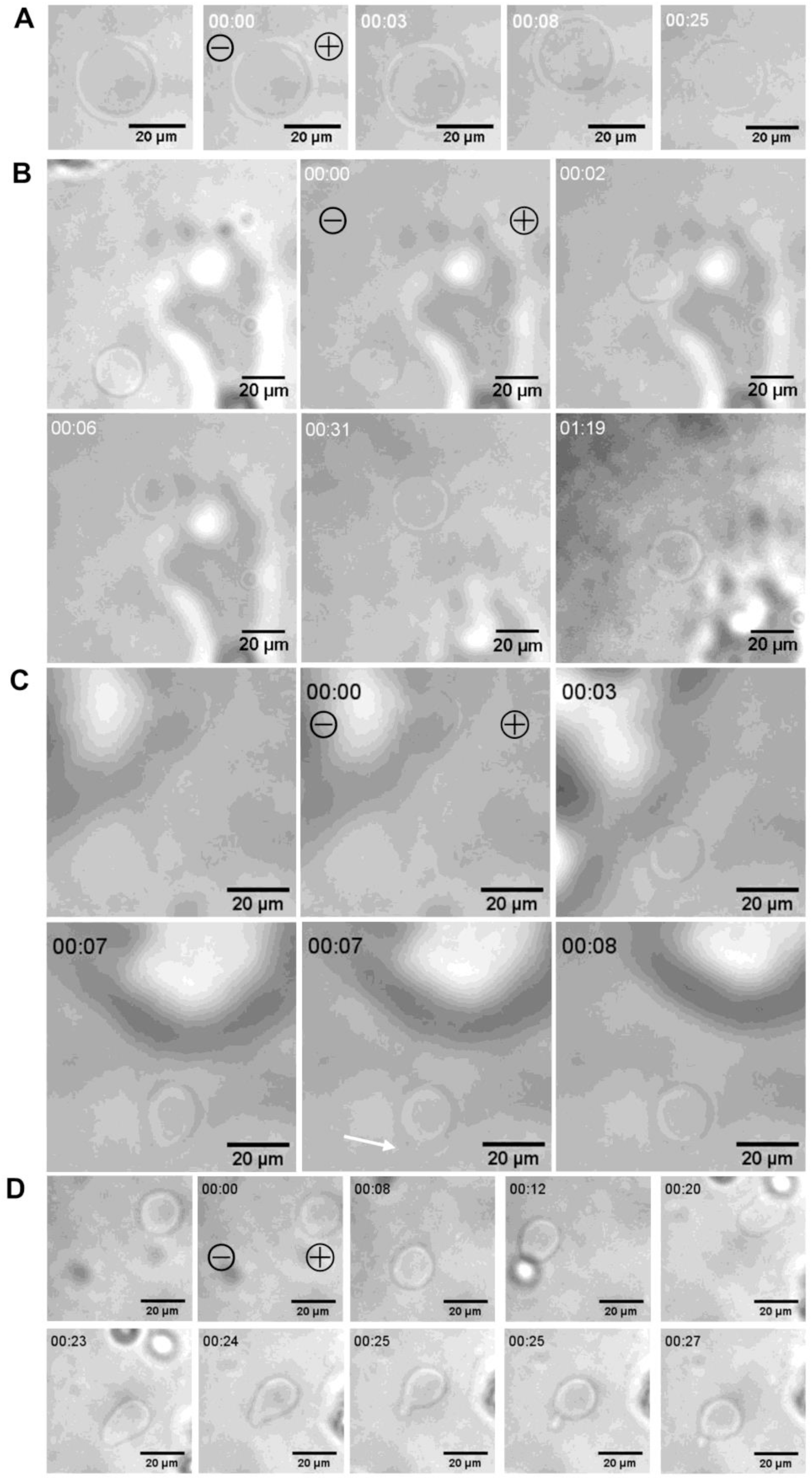
Reactions of several liposome types with 1.1 kV/cm, 400 μs single PEF. A) Lecithin-only liposomes did not divide. B) Lecithin/cholesterol = 2:1 (molar ratio) liposomes did not divide. C) Lecithin/cholesterol = 1:1 (molar ratio) liposomes budded daughter liposomes. The white arrow indicates the budding position. (D) Lecithin/cholesterol = 1:1.5 (molar ratio) budded daughter liposomes. All time displays are in seconds. “00:00” = timing of PEF application. ⊕ = the anode side of the electrode; ⊖ = the cathode side of the electrode.

PC is a main component of lecithin. Previous studies have used DOPC, POPC, and 1,2-dipalmitoyl-sn-glycero-3-phosphocholine (DPPC), all of which are classified as PC. To evaluating which elements contributed to the liposome division, we prepared the liposomes, with each lipid and cholesterol at a 1:1 molar proportion. DOPC produced lecithin-like liposomes, but POPC did not (Fig. 3). Because POPC had higher phase transition temperature than DOPC and temperature influences liposome production [36], we anticipated that the surface aggregation on POPC liposomes may be aggregates of gel-phase POPC. We did not prepare DPPC liposomes because DPPC has a much higher phase transition temperature, making it unlikely for DPPC liposomes to produce lecithin-like liposomes. No division occurred in DOPC liposomes, which most likely resembled lecithin liposomes (Fig. 3A), even though the cholesterol fraction increased by more than 50% (Fig. 3B and C). These results suggest that several lipids together induced division, rather than only one element of phospholipids contributing to the division.

**Figure 3.**
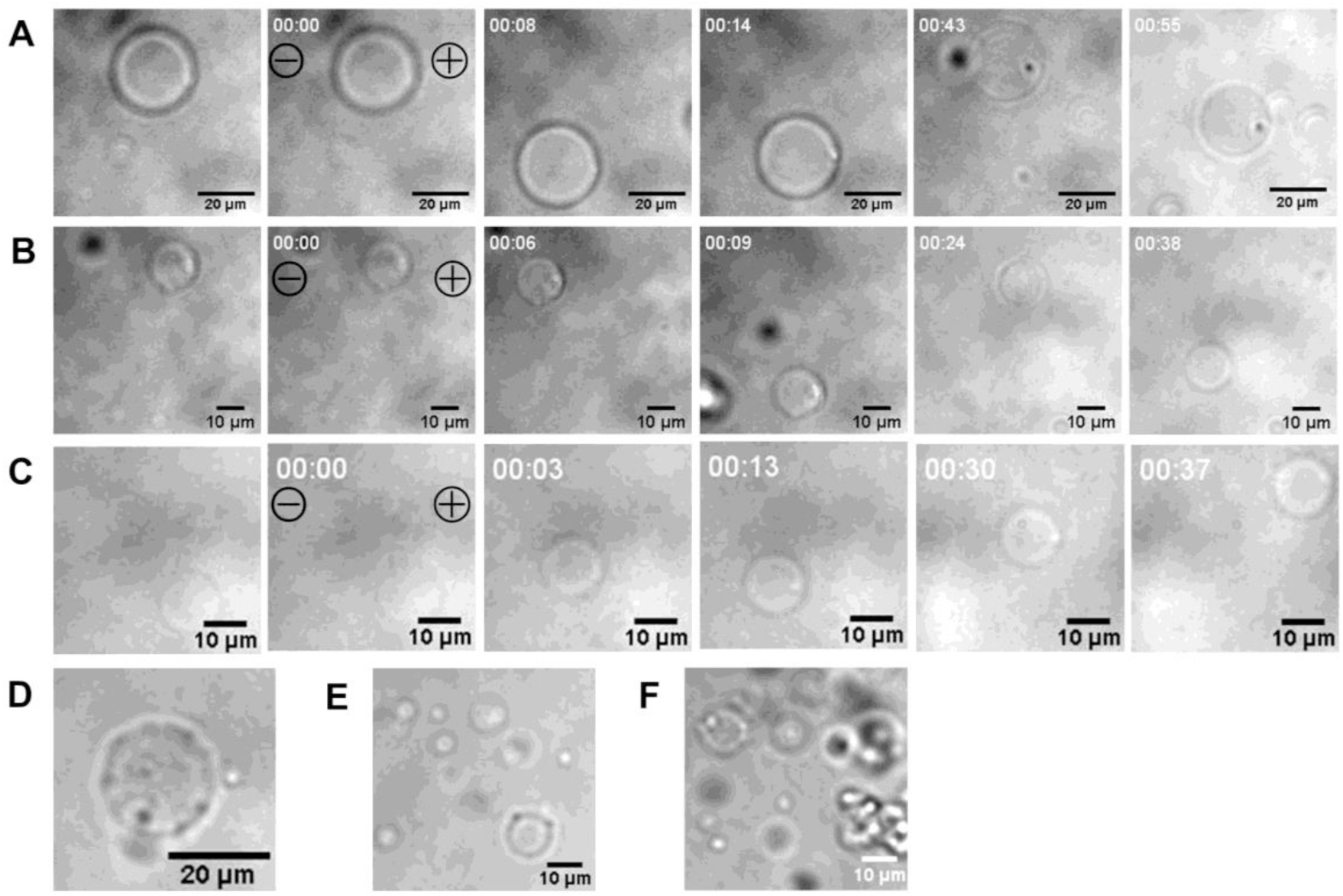
Liposomes without lecithin did not exhibit division. PEF was 1.1 kV/cm, 400 μs single pulse. A) DOPC/cholesterol = 1: 1 (molar ratio). B) DOPC/cholesterol =1:2 (molar ratio). C) DOPC/cholesterol = 1:3 (molar ratio). D) POPC-only liposome. E and F) POPC/cholesterol = 1:1 (molar ratio). All time displays are in seconds. “00:00” = timing of PEF application. ⊕ = the anode side of the electrode; ⊖ = the cathode side of the electrode.

### Conductivity of liquids related to the division

When we lowered the conductivity of both inner and outer liquids to 105 μS/m, no division occurred (lecithin/cholesterol = 1:1) (Table. 3). Because the emulsion method required heavier inner liquids than outer liquids, we mixed glucose solution and PBS (-) for the inner liquid to ensure that inner conductivity decreased to 0.6 S/m. For the same reason, we did not execute the experiments with 1.6 S/m inner liquid. Instead, 0.6 S/m for the inner liquid and 1.6 S/m for the outer liquid (the same as for the red blood cells in PBS) were deemed appropriate.

**Table 3.** Several conductivity patterns of inner and outer liquids. PEF was 1.1 kV/cm (400 μs). σ_ex_ is the conductivity of the outer liquid, whereas σ_in_ is the conductivity of the inner liquid.

| Condition | Division |
| --- | --- |
| $\sigma_{\text{ex}} = 1.6 \text{ S/m}$, $\sigma_{\text{in}} = 0.6 \text{ S/m}$ | $\circ$ |
| $\sigma_{\text{ex}} = 105 \mu\text{S/m}$, $\sigma_{\text{in}} = 0.6 \text{ S/m}$ | × |
| $\sigma_{\text{ex}} = 1.6 \text{ S/m}$, $\sigma_{\text{in}} = 105 \mu\text{S/m}$ | × |
| $\sigma_{\text{ex}} = 105 \mu\text{S/m}$, $\sigma_{\text{in}} = 105 \mu\text{S/m}$ | × |

$\circ$ indicates liposome division and × indicates no liposome division. For $\sigma_{\text{in}} = 105 \mu\text{S/m}$ , we used sucrose solution and for $\sigma_{\text{ex}} = 105 \mu\text{S/m}$ , we used glucose solution.

### Ca-ion influx had little relation to division

To analyze ion influx under the PEF that induced the liposome division, we attempted to observe Ca-ion influx into liposomes. However, we detected no Ca-ion flow into liposomes after a single PEF (Fig. 4). Lecithin liposomes, the surfaces of which are not smooth, collapsed immediately after PEF exposure (Supplementary Fig. 1A). When the liposomes disintegrated, green fluorescence emerged from the outside of the liposomes, indicating that Fluo-8 in the liposomes had seeped to the surroundings and connected to Ca ions (Supplementary Fig. 1B). This proved that our method could detect Ca-ion influx. Because a Ca ion is much smaller than dyes such as propidium iodide, Ca ions are prone to flowing across lipid membranes faster than dyes [37]. Our failure to detect a Ca influx indicated that the extent of liquid flow accompanied by the single 1.1 kV/cm, 400 μs PEF was small.

**Figure 4.**
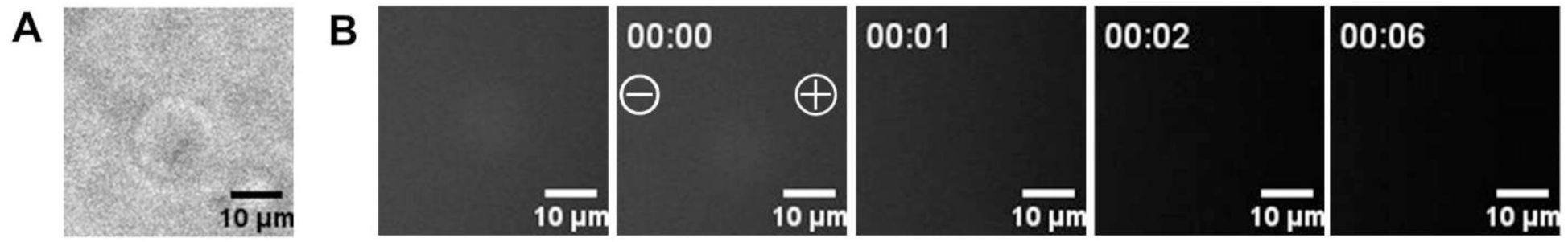
Calcium ion influx was not detected in lecithin/cholesterol = 1:1 (molar ratio) liposomes with 1.1 kV/cm, 400 μs single PEF. A) Bright-field photo of liposomes before PEF application. B) Fluorescent photos of liposomes with a GFP filter. There was no signal after pulse application. All time displays are in seconds. “00:00” = timing of PEF application. ⊕ =the anode side of the electrode; ⊖ = the cathode side of the electrode.

### Endocytosis-like phenomena are also induced

Lecithin-cholesterol liposomes sometimes showed that lipid patches were detached from the membrane surface as in endocytosis (Fig. 5A). Additionally, lipids aggregated on the membrane just after PEF application (Fig. 5B). However, on DOPC-cholesterol liposomes, which did not have division ability, lipid aggregation and patch uptake did not occur (Fig. 5 C, D).

**Figure 5.**
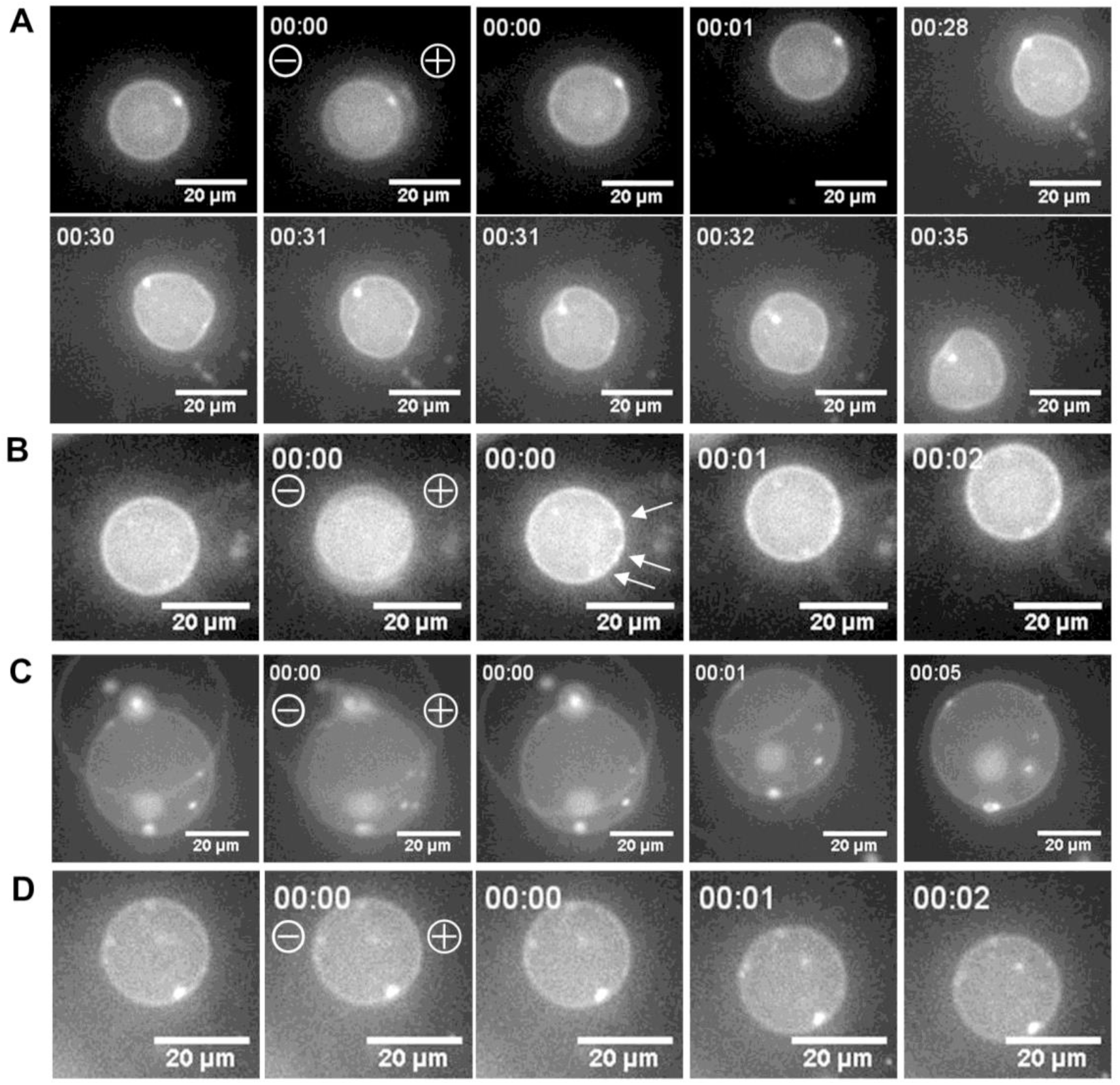
(A) Lecithin/cholesterol = 1:1 (molar ratio) liposomes sometimes exhibited endocytosis-like behavior after a single 1.1 kV/cm, 400 μs PEF application. B) Lecithin/cholesterol = 1:1:1 (molar ratio) liposomes showed patch-like aggregation of lipids on the membrane just after PEF application. White arrows indicate aggregated points. C and D) DOPC/cholesterol = 1:1 (molar ratio) liposomes did not show patch-like aggregation of lipids on the membrane. All time displays are in seconds. “00:00” = timing of PEF application. ⊕ = the anode side of the electrode; ⊖ = the cathode side of the electrode.

## Discussion

### Why did liposomes divide?

Why current conditions trigger liposome division and endocytosis-like behavior remains unknown. Previous studies have reported that electrofusion of liposomes promotes spontaneous liposome division, explaining the phenomenon in terms of thermodynamics; as the amount of lipid per liposome increased by electrofusion, the system became unstable, and consequently, liposomes divided to increase entropy and stabilize the system [38,39]. In our case, however, electrofusion did not precede division, so the previous theory did not explain our results. Because liposome division accompanied liposome deformation, it seemed possible to describe the phenomenon in terms of liposome volume change, but the influence of volume change is likely small because the osmotic pressures of the inner and outer liquids were almost identical. In fact, calcium-ion influx was too small to detect.

In addition, we have no reasonable explanation on why liposome divisions did not occur in previous studies. However, our study with less-conductive inner and outer liquids suggests that liquid conductivity is one of the determining factors. Indeed, one paper using buffered solution as an electroformation buffer showed vesicle budding without releasing daughter vesicle [40].

### Membrane conditions may be related to liposome division

We increased Ca-ion influx by reducing the osmotic pressure of the outer liquid compared with the inner liquid. Using 154 mM glucose solution with a conductivity of 0.7 S/m and a theoretical osmotic pressure of 154 mOsm/L as the outer liquid and PBS(-)-glucose solution with a conductivity of 0.6 S/m and a theoretical osmotic pressure of 308 mOsm/L as the inner liquid, Ca-ion influx was not detected in lecithin liposomes (Supplementary Fig. 2A and B), whereas lecithin-cholesterol liposomes disintegrated with Ca-ion influx (Supplementary Fig. 2C and D). These tendencies coincided with those of previous studies that revealed that liposomes, including charged lipids, such as cholesterol and POPG, were likely to collapse more easily due to PEF exposure compared with those without charged lipids [41,42]. This prompted the question of whether charged lipids can trigger liposome division. We therefore examined the behavior of lecithin-POPG liposomes and lecithin-POPC liposomes. POPG has a negative charge, whereas POPC does not, but both have similar hydrophobic backbones with one saturated and one unsaturated fatty acid chain. Both POPG-containing liposomes and POPC-containing liposomes divided (Supplementary Fig. 2E and F). This suggested that it is not lipid charge but membrane condition that induces liposome division. In fact, Riske et al. reported changes in lipid composition in a plasma membrane during mitosis [43]. Determining which type of lipid can induce liposome division with lecithin should be subjected to a screening wand, and we intend to follow this line of inquiry in the future.

### Future tasks

There are several important points to examine in the future: the relationship between PEF conditions and the time lag from PEF application to division or endocytosis, the influence of inner-liquid viscosity, the relationship between PEF conditions and the size of mother liposomes and that of daughter liposomes, PEF direction and division or endocytosis polarities, and how membrane composition or conditions affect the division.

## Acknowledgments

This study was partially supported by a Grant-in-Aid for Scientific Research (17H03220). The authors would like to thank Enago (www.enago.jp) for the English language review.

## Author contributions

G.U, M.S, and T.O contributed to the experiments. S.K supervised and managed this project.

## Supplementary Figures

**Supplementary Figure 1.**
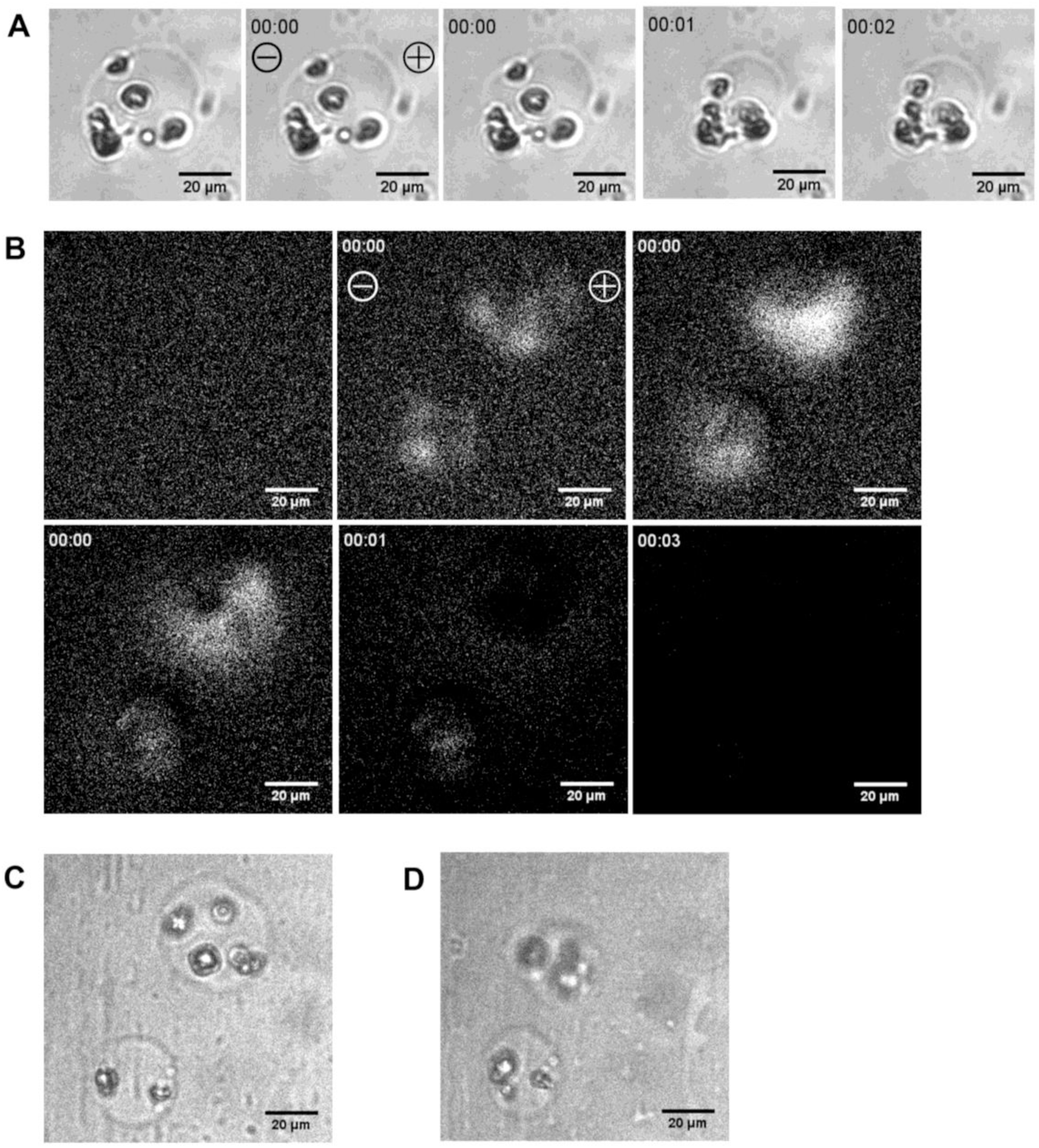
A) Lecithin/cholesterol = 1:1 (molar ratio) liposomes with a rough membrane surface collapsed just after application of a single 1.1 kV/cm, 400 μs PEF. B) Just after integration of lecithin/cholesterol = 1:1 (molar ratio) liposomes, a fluorescent signal emerged within 1 s. C) Bright-field photo of the liposomes before PEF application. D) Bright-field photo of the liposomes after PEF application. All time displays are in seconds. “00:00” = timing of PEF application. ⊕ = the anode side of the electrode; ⊖ = the cathode side of the electrode.

**Supplementary Figure 2.**
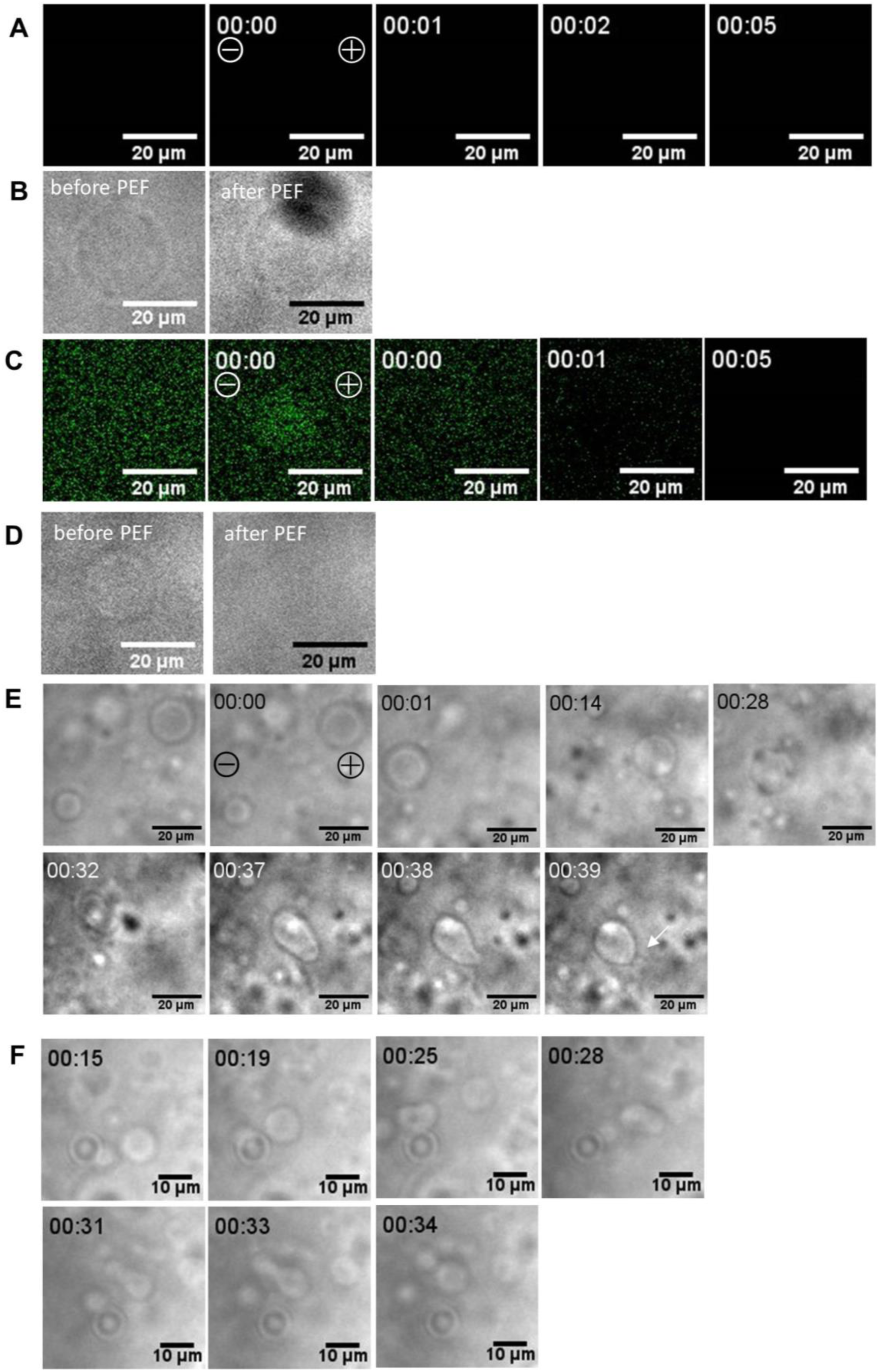
A) Lecithin-only liposomes did not exhibit calcium influx under low-osmotic-pressure outer liquid. B) Lecithin-only liposomes remained after PEF exposure. C) Lecithin/cholesterol = 1:1 (molar ratio) liposomes showed Ca-ion influx just after PEF application under low-osmotic-pressure outer liquid. D) Lecithin/cholesterol = 1:1 (molar ratio) liposomes disappeared after PEF exposure. E) Lecithin/POPG = 2:1 (molar ratio) liposomes divided. White arrow indicates the daughter liposome. F) Lecithin/POPC = 2:1 (molar ratio) liposomes divided. All time displays are in seconds. “00:00” = timing of PEF application.

